# Proteome-scale discovery of protein interactions with residue-level resolution using sequence coevolution

**DOI:** 10.1101/791293

**Authors:** Anna G. Green, Hadeer Elhabashy, Kelly P. Brock, Rohan Maddamsetti, Oliver Kohlbacher, Debora S. Marks

**Affiliations:** Department of Systems Biology, Harvard Medical School, Boston, MA 02115, USA; Biomolecular Interactions, Max Planck Institute for Developmental Biology, 72076 Tübingen, Germany; Department of Computer Science, University of Tübingen, WSI/ZBIT, Sand 14, 72076 Tübingen, Germany; Quantitative Biology Center, University of Tübingen, Auf der Morgenstelle 8, 72076 Tübingen, Germany; Institute for Translational Bioinformatics, University Hospital Tübingen, Sand 14, 72076 Tübingen, Germany; Broad Institute of Harvard and MIT; Institute for Bioinformatics and Medical Informatics, University of Tübingen, Sand 14,72076, Tübingen

## Abstract

The majority of protein interactions in most organisms are unknown, and experimental methods for determining protein interactions can yield divergent results. Here we use an orthogonal, purely computational method based on sequence coevolution to discover protein interactions at large scale. In the model organism *Escherichia coli*, 53% of protein pairs in the proteome are eligible for our method given currently available sequenced genomes. When assaying the entire cell envelope proteome, which is understudied due to experimental challenges, we found 620 likely interactions and their predicted structures, increasing the space of known interactions by 529. Our results show that genomic sequencing data can be used to predict and resolve protein interactions to atomic resolution at large scale. Predictions and code are freely available at https://marks.hms.harvard.edu/ecolicomplex and https://github.com/debbiemarkslab/EVcouplings

## Introduction

A longstanding goal of molecular and structural biology is to determine which proteins interact with each other at residue resolution. Experimental methods have given the research community a rich characterization of many molecular interactions^1–3^, but the vast majority of all interactions are undiscovered or not structurally solved. Estimates extrapolated from yeast two-hybrid studies^3^ suggest that in the model organism *Escherichia coli*, about one in 1,000 of all possible protein pairs interact (10^4^ / 10^7^). Less than 40% of the protein interactions in *E. coli* have been experimentally observed^3^ and at most 4% have been structurally characterized, even in related species. Since these experimental methods are labor- and cost-intensive, there has been increased interest in computational approaches, alone or in combination with experiments. Homology modeling, a computational approach that exploits one known interaction to infer the structure of a similar interaction between homologous pairs, fills some of the knowledge gap^4^, but the challenge remains for *de novo* prediction from sequence alone when homologous pairs of structures are unavailable.

Recently, approaches such as EVcouplings have successfully leveraged the vast corpus of natural sequences to determine 3D structures, by using probabilistic graphical models to infer coevolved positions. These coupled positions can be sufficient to fold single proteins^5–7^ and RNAs^8^, without the use of homologous structures, and to predict protein interactions *de novo* ^9,10^. Ideally, this approach would extend to determine interactions between any pair of proteins, but previous work has been limited to predicting inter-protein contacts between co-operonic proteins, and therefore was calibrated on and usable only for select, small datasets^9–11^. In this work, we solve the challenges of concatenation and statistical scoring to provide a new method that integrates information from residue-level scores to predict protein interaction **(Fig. 1)**. Residue-level coevolution is then used as restraints for molecular docking to provide atomic-resolution models of protein interactions. As a case study, we highlight 620 interactions involving cell envelope proteins, which are some of the most challenging to determine experimentally, and demonstrate the applicability of our method to 53% of the *E. coli* proteome as of 2019.

**Figure 1:**
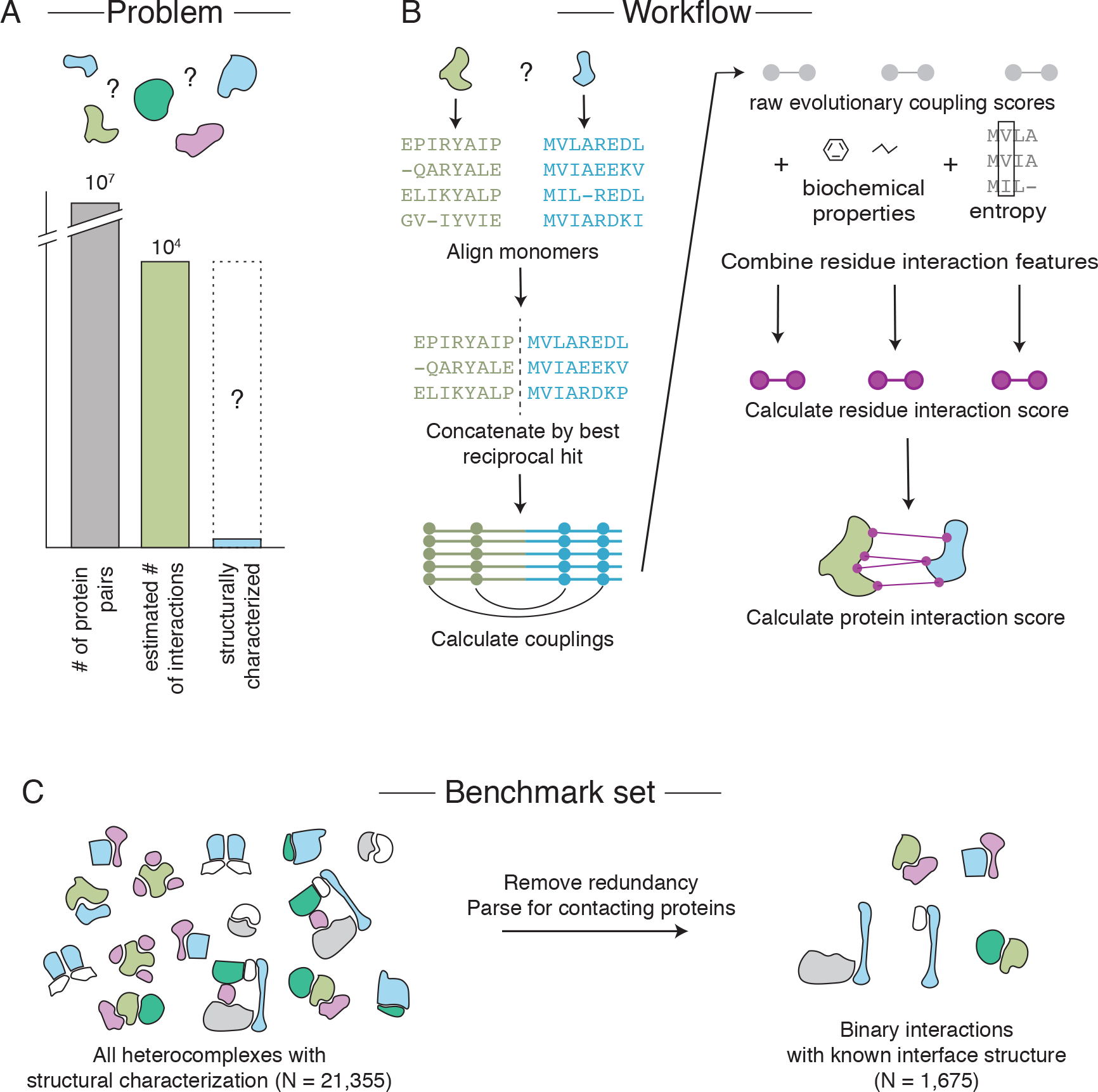
Scaling EVcouplings methods to full bacterial genomes. (A) The search problem for binary protein-protein interactions in *Escherichia coli* involves finding all of the estimated 10^4^ true interactions out of 10^7^ possible pairs. Only approximately 300 of these true direct interactions have structural resolution in any organism. (B) Evolutionary couplings learned directly from protein sequences can resolve interfaces. Sequence alignments of both monomeric proteins are created and concatenated by the best reciprocal hit procedure before inference of evolutionary couplings. Raw evolutionary coupling scores can be combined with features of their distribution, biochemical properties, and sequence entropy to improve inference. (C) A benchmark dataset of all non-redundant protein interactions with known interface structure.

## Results

### *E. coli* has many undiscovered protein interactions that are amenable to EVcouplings

Most protein interactions remain undiscovered or structurally unresolved, even for well-studied model organisms such as *E. coli.* Of the estimated 10,000 protein interactions in *E. coli*, approximately 3,946 have been observed experimentally^3^, involving about 50% of *E. coli* proteins, hence leaving a large fraction of interactions unknown (**Fig. 1**). The majority of the protein monomers in the *E. coli* proteome (3,186 out of a total of 4,391) have high-quality monomer alignments and are therefore amenable to EVcouplings analysis (Methods). For the 67% that have an experimental monomer structure (either in *E. coli* or for a homologous sequence from another organism), 78% have reasonable precision of the top ECs (60% for the top L/2 ECs, where L is the protein length) (**Supplementary Table 1, Supplementary Fig. 1**). We therefore restrict our computational predictions of interactions to the set of 3,186 (*i.e.,* 75% of proteins, covering 53% of the total possible interaction space).

### Computing coevolution across proteins

This work addresses two main challenges in determining large-scale interactions across a whole genome. First, to avoid the use of operonic structure and to test *any* protein pair, we constructed alignments of pairs of proteins from different organisms by identifying the nearest orthologs using reciprocal best hits^12^ (Methods). While the “reciprocal best hit” approach has many assumptions, it errs on the conservative side – we are more likely to get false negatives than positives – as inter-protein coupling scores are lower when pairings are incorrect. Second, the challenge of identifying true interactions is compounded by their relative sparsity compared to all possible pairs – an estimated 0.1% of all pairs of proteins have a direct interaction (**Fig. 1**). Therefore, even small false positive rates will result in large numbers of predicted interactions that are false.

To minimize the number of false positive interactions, we trained and calibrated on both positive and negative benchmarks: a ‘gold standard’ non-redundant set of 560 interactions with known structure, and a set of 2,500 non-redundant “non-interacting” protein pairs with no signal for interaction in yeast two-hybrid experiments (**Supplementary Table 2**, Methods). Evolutionary couplings (ECs) computed across these 560 pairs can recall correct interactions, at the expense of a large number of false positives from the set of 2,500 non-interacting protein pairs (**Fig. 2A**). The accuracy of the model increases when we include other structure-agnostic features in a logistic regression: EC score, sum of intra-protein EC coupling constraints, rank of inter-EC relative to intra-EC pairs, sequence conservation, and hydrophilicity (**Supplementary Table 3,** Methods). Specifically, if we consider a raw EC score threshold that limits our false positive rate to 0.1%, we recover only 12.5% of interacting complexes in the benchmark set. By contrast, the simple logistic regression score achieves a recall of 21% of true interacting complexes with a false positive rate of 0.1%. When the structure of at least one of the monomers is known, the logistic regression model achieves a recall of 26% after incorporation of accessible surface area and precision of intra-molecular evolutionary couplings as features (**Supplementary Fig. 2A**). Our model outperforms yeast two hybrid experiments, achieving a recall of 35% versus the 29% obtained experimentally at a false positive rate of 1%^3^.

**Figure 2:**
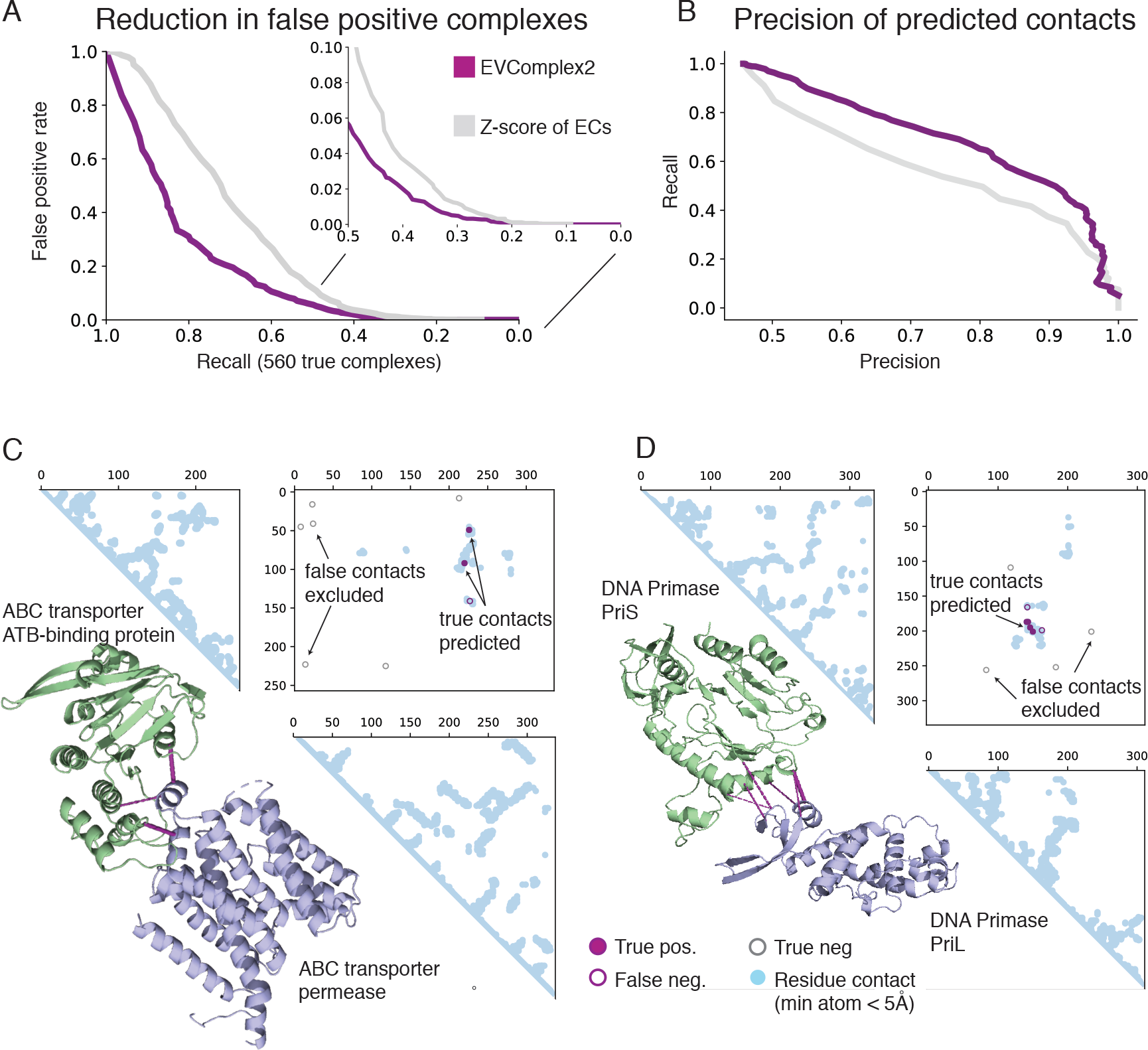
Model predicts interacting residues and interacting proteins with high precision. (A) Our logistic regression model (purple) reduces the false positive rate compared to raw ECs (grey). The x-axis shows the recall on the positive benchmark set of 560 true complexes, and the y-axis shows the false positive rate on the held-out dataset of non-interacting complexes at the score threshold that gives each recall value. (B) Once the interaction of the proteins is known, our model has higher recall of interacting residues than raw ECs. For protein complexes with inferred interaction at a 40% recall cutoff (N=224), the precision versus recall of the top 10 ECs is plotted. ECs are considered true if inter-residue minimum atom > 8Å, and false if > 12Å. (C) Example performance on known interaction between and ABC transporter permease and ATP binding subunit (UniProt IDs Y1470_HAEIN and Y1471_HAEIN, PDB ID: 2NQ2 chains C and A. (D) Example performance on known interaction between and DNA primase PriS and PriL (UniProt IDs PRIS_SULSO and PRIL_SULSO, PDB ID: 1ZT2 chain A and B)

The logistic regression model also outperforms raw ECs at detecting residue contacts when the interaction of the protein pairs is presumed known. Considering only the protein pairs in the positive benchmark set, at a score threshold that gives a precision of 80% of true inter-protein ECs, we recover 54% of the true ECs in our dataset using just raw EC scores, 65% of true ECs with the logistic regression model, and 78% with the logistic regression model when incorporating features of known monomer structures (**Fig. 2D**, **Supplementary Fig. 2B**).

### Prediction of membrane protein interactions

Since the cell membrane contains many protein interactions essential for life, but is notoriously difficult to study experimentally^13^, we targeted the cell envelope proteome for detailed analysis. We based our analysis on 1,583 proteins previously described as localized to the *E. coli* cell envelope^1^, constituting ~1.25 million possible interacting pairs. We assayed each compartment of the *E. coli* cell envelope proteome with itself and with adjacent compartments (Methods) (**Fig. 3A**), for a total of 939,159 protein pairs. After monomer sequence alignment, concatenation, and EC calculation, 191,999 proteins pairs comprised of 1,053 proteins (566 non-redundant protein families) pass quality thresholds for analysis. The majority (771) of these proteins are inner membrane proteins. 49% of these proteins have at least a partial structure known of itself or a homologous protein.

**Figure 3:**
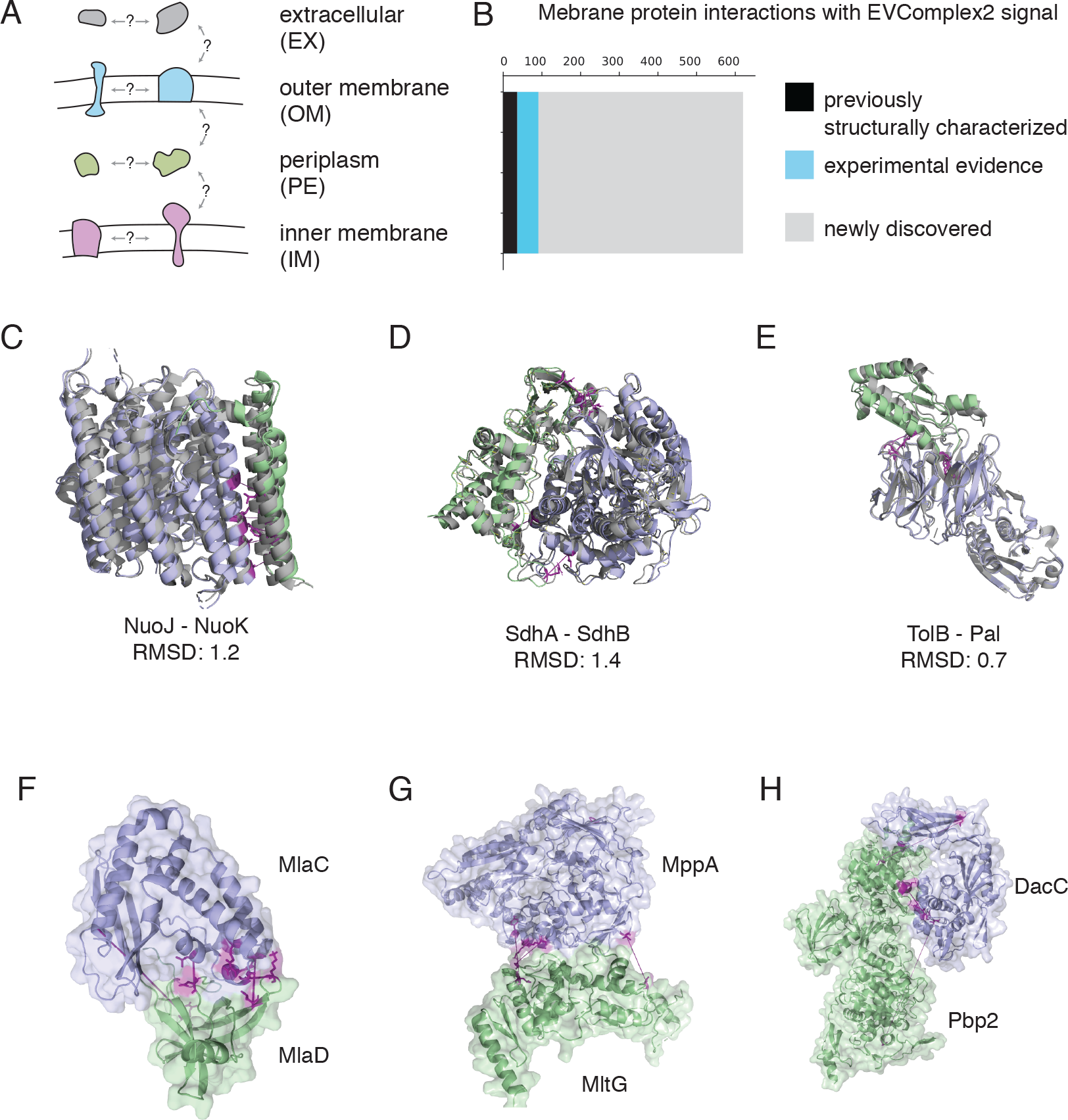
Discovery of hundreds of new interactions in the *E. coli* membrane proteome. We searched a high-value subset of the 10^7^ possible interactions in the *E. coli* proteome by searching membrane compartments with themselves and with adjacent membrane compartments. We found 620 high-scoring protein interactions in the cell envelope, including 29 with structural characterization and 92 with previous experimental evidence. (C-E) 3D configurations of previously structurally characterized interactions accurately predicted by molecular docking with inferred restraints. Known structures are shown in grey. RMSDs calculated by comparison to known structures with PDB IDs 3RKO, 2WU2, and 2HQS, respectively. (F-H) Example of three docked models of newly resolved protein complexes: MlaC/MlaD, MppA/MltG, and DacC/Pbp2.

We predict 529 not previously known protein interactions in the *E. coli* cell envelope as well as the details of their interacting residues (**Fig. 3B**, **Supplementary Table 4, Supplementary Fig. 3**). Despite having chosen a stringent false positive rate threshold of 0.1%, we expect a minority (30%) of these interactions may be false positives due to the large size of the tested space. Indeed, in our negative assessment dataset of 12,333 protein pairs in the membrane unlikely to interact, we detect interaction for 57 (0.5%). Of 171 independent families of protein interactions with known interaction and known crystal structure that can be analyzed by our method, we recover interaction signal for 29 of them, consistent with recall at this threshold from our calibrations.

For protein interactions that have previously been observed experimentally, we report significant residue-level coevolution for 450 pairs (at least 3 residue pairs above the 80% precision threshold in calibration). These predictions can be used to disambiguate directly interacting proteins in complexes identified via AP-MS, such as the Tol-Pal complex reported below. 91 of these protein pairs would have been blindly identified as interacting by our stringent scoring system.

The protein interactions in the *E. coli* membrane with the highest interaction score are involved in essential cellular processes; for instance, the RodA and penicillin-binding protein cell wall polymerases, which we had previously characterized in a detailed study^14^, as well as components of the electron transport chain, components of the ATP synthase, and components of the bacterial flagellum.

### Atomic resolution models of membrane protein complexes

The inter-protein residue pairs identified by our method can be used as restraints for molecular docking, to resolve the 3D structure of protein complexes^9,10^, (**Fig. 3C-E).** Here, we provide detailed resolution and characterization of newly structurally resolved protein complexes that score highly in our interaction model.

We provide structural models for high-scoring, previously structurally uncharacterized interacting pairs. We also resolve the MlaC–MlaD proteins **(Fig. 3F)**, which are involved in inner membrane phospholipid transport, and whose interaction was previously observed experimentally^1^. We resolve the interface for the newly discovered interaction between MppA and MltG **(Fig. 3G)**, which are involved in murein synthesis in the cell wall, and between the peptidoglycan biosynthesis protein DacC and the peptidoglycan polymerase Pbp2 **(Fig. 3H)**. While this docked model only satisfies one of five inter-molecular restraints, it is possible that movement by the Pbp2 protein during synthesis may change the configuration of the interface.

Our method resolves the interaction interface between the flagellar proteins FlgL and FlgK, which form the junction between the flagellar filament and flagellar hook^15^. Extrapolating from our docked model of the monomers and a previously produced model of the FlgK ring from *Campylobacter jejuni,* we infer the configuration of an 11-mer ring of FlgL inside an 11-mer ring of FlgK^16^ **(Fig. 4**, **Supplementary Fig. 4)**.

**Figure 4:**
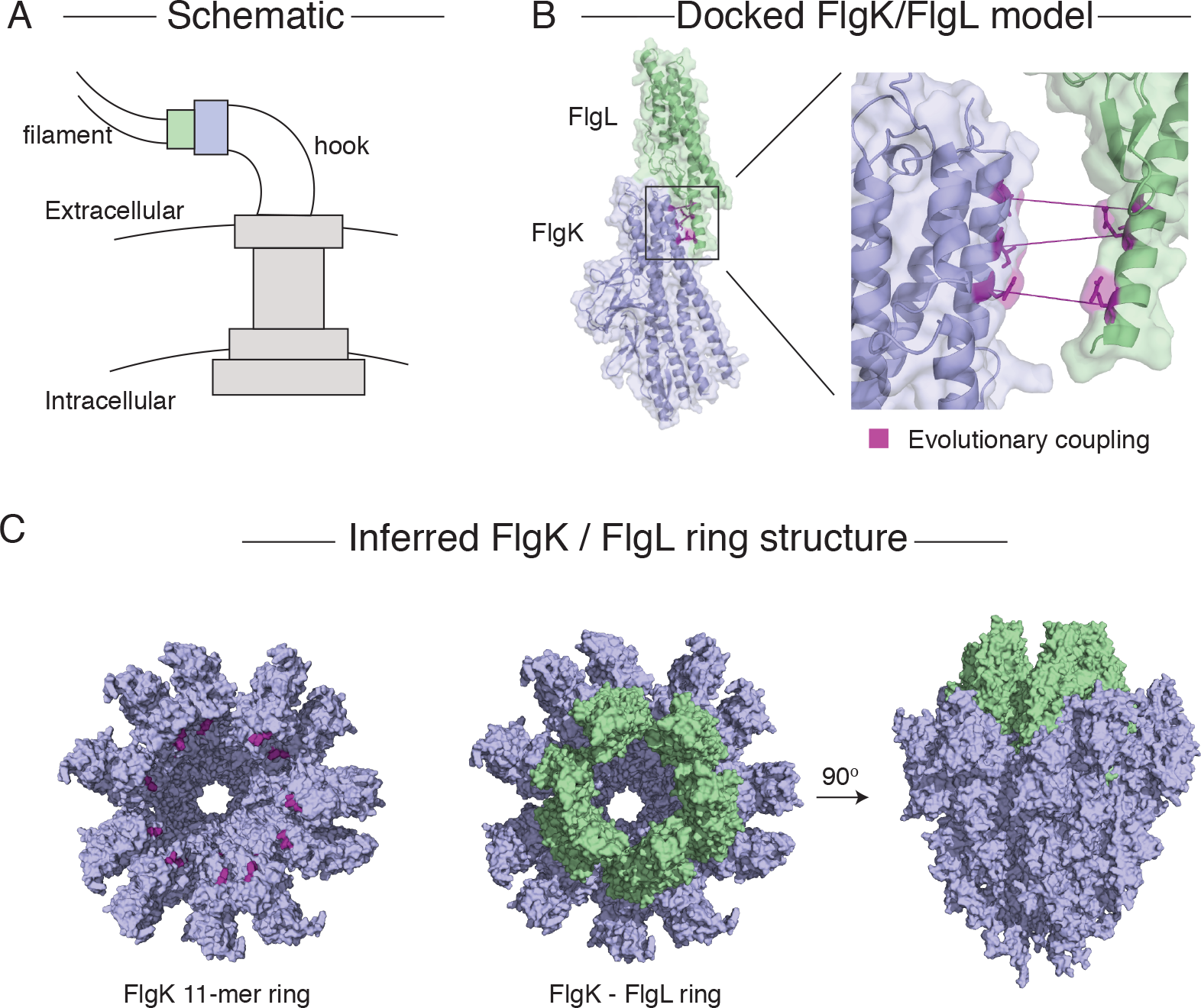
Model of the flagellar hook-filament junction. (A) Schematic of the orientation of the bacterial flagellum. The proteins FlgL (green) and FlgK (blue) form two rings which create the junction between the hook and filament of the flagellum^16^. (B) Docked model of FlgL and FlgK using evolutionary couplings. PDB structures of homologous proteins from *Salmonella typhimurium* were used in docking (PDB IDs 2D4Y and 2D4X, respectively). Predicted interface residues are highlighted in purple. (C) A previously inferred model of the FlgK ring from *Campylobacter jejuni^16^* was used to infer the structure of the entire hook-filament junction. Evolutionary coupled residues (purple) show the interface for FlgL ring insertion into the FlgK ring. By aligning our docked model to the *C. jejuni* ring, we show that an 11-mer ring of FlgL fits inside the FlgK structure.

We constructed an atomic model of the Tol/Pal system (**Fig. 5**), a protein machine that spans the inner and outer membranes of Gram-negative bacteria. These proteins may play a role in membrane constriction during bacterial cell division^17,18^. For the known interacting interface between TolB and Pal^19^, we accurately recapitulate the known structure (PDB ID 2HQS). We predict interactions and residue contacts for the known interaction between TolA and TolB^3,20^ as well as for the interactions of TolA, TolR, and TolQ in the membrane^21^. The protein CpoB interacts with the Tol system and the cell division septum site^18^, and here we provide a molecular model of its interaction with TolB.

**Figure 5:**
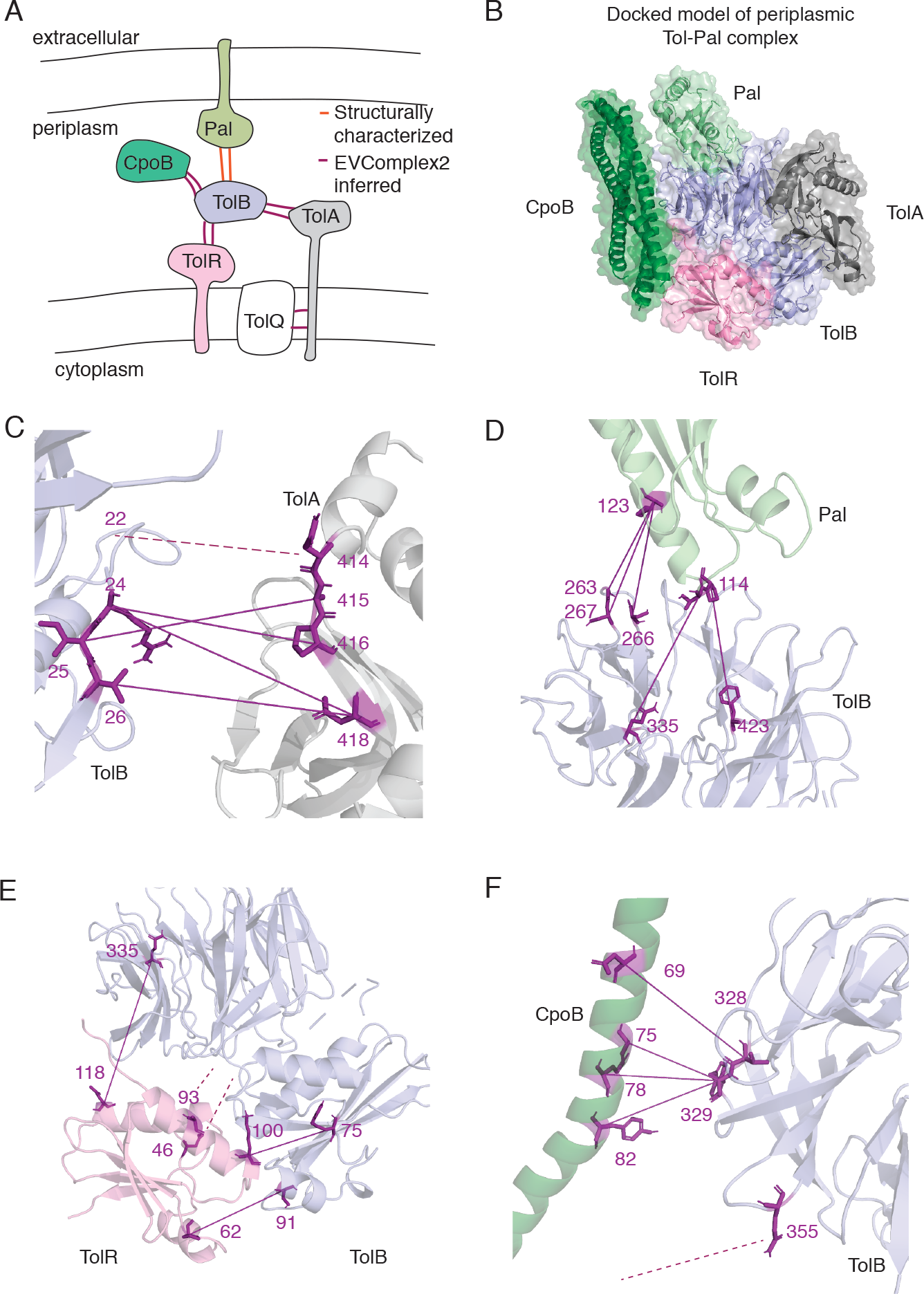
Atomic resolution model of the Tol-Pal complex produced by EVcouplings. (A) Schematic of the proteins involved in the Tol-Pal complex1 (TolABRQ, CpoB, and Pal). Interactions with previously solved interfaces are shown in orange and interactions inferred by our method are shown in purple. (B-E) Residue resolution of TolB-Pal, TolB-TolA, TolR-TolB, and CpoB-TolB interfaces. The top 5 inferred interface contacts are shown in purple. Dashed lines indicate inferred contacts where one or more residues is missing from the solved structure. Structures used are 1TOL_A (TolA), 2HQS_A (TolB), 5BY4_A (TolR), 2HQS_H (Pal), and 2XDJ_A (CpoB). (F) Complete model of the Tol-Pal complex inferred by aligning results of docked pairwise models. Note that CpoB is inferred to be a trimer *in vivo* but was docked as a monomer for modeling purposes.

## Discussion

In this work, we demonstrate a method for inferring protein interactions at residue-level detail from genomic sequences alone. We provide a probabilistic scoring method, calibrated on large, non-redundant datasets, which can be used to determine the probability of interaction of two proteins. If the protein interaction is known, or inferred by the protein interaction score, the residue interaction score can be used directly to determine residues in contact at the interface. These methods can be freely accessed via our webserver or by downloading the source code ^22^ (https://github.com/debbiemarkslab/EVcouplings).

Our method assumes that protein interactions are conserved across species, an assertion which is certainly not true for all complexes and will lead to false negatives in the case of newly evolved or poorly conserved interactions. In addition, any method to predict physical contacts in protein interactions must be calibrated on known structures, and therefore may inherit biases from the set of proteins that have been structurally characterized. In particular, crystal structures are known to be biased against membrane proteins and transient interactions, which could lead to false negative predictions.

With current sequence databases, we are able to apply this method to 53% of the protein pairs in the *E. coli* proteome. By using our best reciprocal criterion, we limit ourselves to one ortholog per genome, which limits the amount of sequence diversity that can be found for families with multiple paralogs per genome. Growth in sequence databases and development of fast concatenation techniques to handle paralogous sequences will increase the number of protein pairs that can be analyzed.

Method development using coevolution to detect protein-protein interactions has thus far been tested on a handful of well-studied protein-protein interactions^11,23–25^, reaffirming our ability to predict well-conserved, well-characterized protein interactions. While there are many possible avenues for continuing to improve coevolution-based prediction of protein interactions, we suggest that future development should begin with large, non-redundant biological datasets such as ours. By identifying and diagnosing cases where existing methods fail on real datasets, new developments will lead to greater opportunities to discover unknown protein interactions.

## Acknowledgements

We thank members of the Marks, Sander, and Kohlbacher labs for discussion and feedback. We thank Dr. Thomas Hopf for input on EVcouplings methodology and Dr. Anastasiya Yakhnina for input about the Tol-Pal complex. Computational resources and support were provided by the Orchestra High Performance Compute Cluster at Harvard Medical School, which is funded by the NIH (NCRR 1S10RR028832-01). AGG was supported by National Science Foundation Graduate Research Fellowship DGE1144152. HE is funded by the International Max Planck Research School (IMPRS) "From Molecules to Organisms.”

## Note

while preparing this manuscript, Cong et al. published complimentary work, DOI:10.1126/science.aaw6718

## Methods

### Creation of positive benchmark set

A list of all heteromeric structures in the PDBePISA was downloaded^26^ (download date: 2/19/2018) and passed through the RCSB Protein Data Bank with the following query: “Sequence Length is between 30 and 1200 and Resolution is between 1.0 and 3.0 and Representative Structures at 30% Sequence Identity.” IDs were mapped to UniProt ^27^ identifiers using the SIFTS database^28^ (download date 2/20/2018). Only complexes with at least two chains that map to different UniProt identifiers were kept. Single-chain and fusion proteins were removed because they may have non-specific interface contacts not found in nature.

For each protein, we extracted the PFAM domains^29^ annotated in that protein, which we call the PFAM set for that protein. We then consider a protein-protein interaction as unique if and only if the interacting proteins constitute a pair of PFAM sets not yet seen in our database, yielding 4,154 non-redundant pairs of interactions. We then consider only contacting pairs, defined as a protein pair with at least 20 pairs of residues with minimum atom distance < 5Å. Because each unique interaction can be represented by many crystal structures, representative crystals were chosen based on the number of other interfaces present in that crystal – *i.e.*, complexes with more subunits were prioritized. We removed any cases where the two proteins map to the same chain, where the two proteins share any PFAM domain, or where either of the proteins was less than 30 amino acids in length, resulting in a final list of 1,675 protein complexes with known interfaces (**Supplementary Table 2)**.

### Creation of negative benchmark set

2,500 negative examples were selected randomly from all pairs of complexes with no signal for interaction in a large-scale Y2H experiment in *E. coli*^3^. We verified that no pair of proteins had the same PFAM set as any other pair of proteins in the negative set, and that no pair had the same PFAM set as any pair of proteins in the positive benchmark set **(Supplementary Table 2)**. For a negative assessment dataset to evaluate the performance of our predictor on the *E. coli* cell envelope, all pairs of proteins where the monomers are found in different AP-MS benchmark complexes were assessed^1^.

### Monomer Sequence Alignment

For alignments of the positive benchmark set, all UniProt identifiers corresponding to the sequence of interest were extracted from PDB^30^. For alignments of the *E. coli* genome, all identifiers from the reviewed *Escherichia coli* (strain K12) reference proteome were downloaded from UniProt^27^. The entire length of every protein was used for sequence alignment. Alignments were constructed using jackhmmer^31^ with five iterations against the UniProt database (downloaded February 2018), as implemented in the EVcouplings software package. Alignments were considered to have sufficient coverage if at least 80% of the columns were less than 50% gaps. A range of bitscores were tested (0.1, 0.2, 0.3, and 0.5, and 0.8 times protein length), and the alignment with the largest N_eff_ that had 80% coverage of columns was selected for each protein. Proteins from the *E. coli* genome were deemed eligible for EVcomplex analysis if their monomer sequence alignment had at least 80% column coverage and N_eff_/L >= 2.5, after characterizing the precision of monomer protein couplings across a range of sequence alignment diversities (**Supplementary Fig. 1**).

### Concatenation

Monomer protein sequences were concatenated by extracting the species annotation from the UniProt database. A single representative for each species was chosen based on best reciprocal identity to the query sequence, after excluding closely related paralogs in the query genome with more than 90% sequence identity. If, for a single species, a best reciprocal hit for both monomers can be identified, those hits are concatenated and added to the alignment.

### Concatenated alignment quality control

To avoid analyzing protein pairs where the two proteins are homologous, we implemented three different quality control metrics. First, we remove all pairs of proteins where the first protein contains a PFAM domain that is found in the second protein. Then, we remove all pairs where our structure comparison protocol found overlapping hits to the same chain of the same PDB structure. Finally, we remove all pairs that display high-scoring coevolution along a diagonal between the two proteins (*i.e,* between position *i* in protein A and the corresponding position *i* in protein B). We consider these contacts to be artifactual due to their very high scores relative to known interactions. In addition, we do not consider proteins from *E. coli* where there is a non-traditional amino acid annotated in the UniProt sequence, because these can indicate pseudogenes.

To ensure high-diversity, low gap alignments for inference, we removed from consideration concatenated sequence alignments with N_eff_/L less than 0.2 or coverage of columns lesser than 80%.

### EC calculation

For EVCouplings calculation, hits composed of more than 50% gaps were filtered from the alignment, and sequences with homologs more than 80% identical were downweighed to compute N_eff_, the effective number of sequences^5^. ECs were calculated using pseudo-likelihood maximization^32,33^, as implemented by Weinreb *et al.* (2016)^8^. The λ_J_term was scaled by the number of amino acids minus one times the number of sites in the model minus one. Pre- and post-processing was performed using the EVCouplings Python package^22^.

### Comparison of ECs to experimental structures

To identify experimental structures for comparison of evolutionary couplings, the monomer sequence alignment were searched against the PDB database using *hmmsearch*^31^. For *E. coli* monomer alignments, the top 20 were selected. For the validation dataset, all hits above a bitscore threshold of 0.2 *L* were selected. Once structures were selected for comparison, the minimum of the minimum atom inter-residue distances across all models were used for comparison to ECs^6^.

### Solvent accessible surface area calculation

Relative and absolute solvent accessibilities were calculated using DSSP^34^ with a probe size of 1.4 Å and van der Waals radii from literature^35^. If multiple structural hits were found for each protein, we take the mean relative solvent accessibility across all structures, to avoid biases caused by incomplete structures.

### Conservation calculation

The conservation *C* for each column *i* in the concatenated sequence alignments was calculated as one minus the normalized entropy for the column *i*,

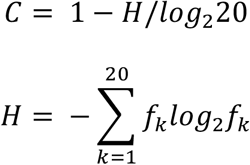

Where *f_k_* is the frequency of the *k*-th amino acid in column *i*.

### Detecting inter-protein residue contacts

For analysis of raw ECs, the Z-score of each inter-residue EC was calculated using the distribution of all inter-residue ECs.

The model for residue interaction was trained on the positive benchmark dataset. Our dataset for includes 560 protein complexes for which the concatenated sequence alignment has sufficient coverage and diversity. Because most of the inter-protein ECs are noise and do not correspond to real contacts^9,10^, we take the top 10 inter-protein ECs as input to the model. This resulted in 5,334 residue pairs with a defined minimum atom distance, 26% of which were within 8Å.

We trained a logistic regression classifier to predict the probability that two residues *i* and *j* are in contact (distance < 8Å), given their EC score and additional residue-level features. 25% of the data was held out for testing. Models were trained using the Logistic Regression classifier in Scikit-learn^36^, using the liblinear solver and L1 regularization. Regularization weight of 0.05 was chosen to maximize a tradeoff between recall and precision using 10-fold cross validation.

The features with non-zero weights were the Z-score of the APC-corrected FN score^5^, the conservation of the more conserved for the two residues, the relative rank of the first inter-protein EC, the maximum of the intra-protein EC enrichment for the two residues^6^, and the combined hydrophilicity score of the amino acids at those two positions.

For a model to be used when monomer structures are known, we repeated the model selection procedure with two additional features: the minimum of the accessible surface area of both residues, and the minimum of the precision of the intra-protein ECs ranked above the current inter-protein EC. These two features were found to be significant in combination with the features found above.

### Detecting protein interactions

To build a model for determining whether two proteins interact, we combine the score of the residue interaction model for the top ten inter-protein residue pairs using a logistic regression framework. The training dataset was composed of positive and negative examples. Positive examples were protein pairs from the positive benchmark set for which at least three of the top 10 inter-protein ECs have distances within 8Å, for a total of 188 complexes. An equal number of negative examples were chosen from the negative benchmark set. 25% of this combined dataset was held out for testing.

We trained two models, using the residue interaction score with and without structure-aware features. We also include the relative rank of the highest-scoring inter-protein EC as a feature. Models were trained using the Logistic Regression classifier in Scikitlearn^36^, using the liblinear solver and L1 regularization (weight = 0.1), chosen to maximize a tradeoff between recall and precision using 10-fold cross validation.

### Determination of false positive rates for interaction detection

The protein pairs from the negative benchmark set that were excluded from the training and test set were used to assess the false positive rates expected when the model is applied to large datasets.

### Cell envelope proteome

Localizations of *E. coli* proteins were extracted from a previous study^1^. Cell envelope proteins were separated into four categories: inner membrane bound (integral or lipoprotein), periplasmic, outer membrane bound (integral or lipoprotein), and extracellular. Proteins annotated as “membrane related” but without a specific location were excluded. Each of the four compartments was tested against itself and against proteins in the physically adjacent compartment(s): inner membrane versus periplasm, periplasm versus outer membrane, and outer membrane versus extracellular.

### Docking

Side chains of all monomers were perturbed and optimized by using SCWRL4^37^ to revoke any effect of crystallization in a complex that may bias the docking procedure. Restraints-based docking was done using HADDOCK (High Ambiguity Driven biomolecular DOCKing) v2.2^38^ and CNS^39^. This allows restricting the possible interaction search space using a more sophisticated treatment of conformational flexibility. Top five inter-ECs scores in both of structure-aware and structure-free regression models were selected. Unambiguous distance restraints were applied on C-beta (except for glycine, where C-alpha was used) with an effective distance of 5 Å with upper and lower bound of 2 Å. Default HADDOCK protocol was done starting with a rigid-body energy minimization (500 models), followed by semi-flexible refinement in torsion angle space (100 models), and ending with further refinement in explicit water (100 models). The explicit flexibility during the molecular dynamics refinement HADDOCK can account for small conformational changes occurring upon binding. The resulting models were scored using the default HADDOCK scoring function, and the top ten best-scoring models, as well as the top models from RMSD clusters, are reported.

