## Supplementary Figures for "Proteome-scale discovery of protein interactions with residue-level resolution using sequence coevolution"

**Supplemental Figure 1: High quality sequence alignment of protein monomers from *E. coli*.** (A) The precision of the top ECs increases as the effective number of sequences ($N_{eff}$) divided by sequence length (L) increases, with a plateau at approximately $N_{eff}/L = 2.5$. Each point corresponds to a monomeric protein in *E. coli* for which there is a crystal structure or structure of known homolog. The X axis was truncated at 20 to show detail in the low range. (B) A histogram of the precision of the top L ECs for proteins whose sequence alignments had $N_{eff}/L$ >= 2.5

A

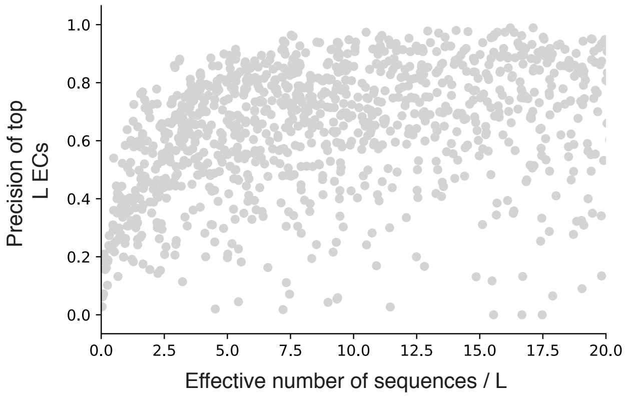


B

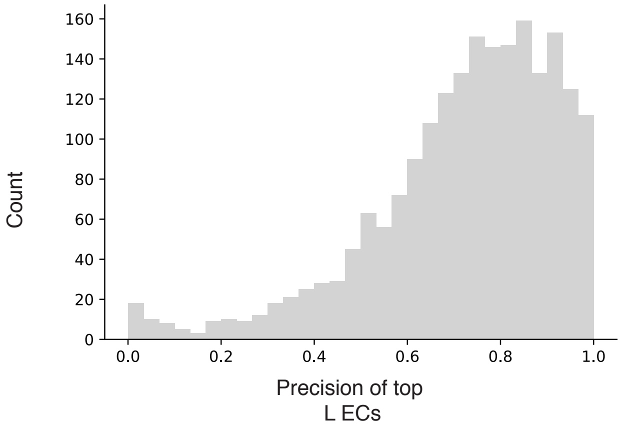

**Supplemental Figure 2: Features of monomer protein structure further increase precision and recall** (A) Our logistic regression model (purple) reduces the false positive rate compared to raw ECs (grey). A logistic regression model that also incorporates features of monomeric protein structure further outperforms the structure-agnositc logistic regression. The x-axis shows the recall on the positive benchmark set of 560 true complexes, and the y-axis shows the false positive rate on the held-out dataset of non-interacting complexes at the score threshold that gives each recall value. (B) Once the interaction of the proteins is known, incorporation of features of monomeric structures greatly increases model recall for a given precision. For protein complexes with inferred interaction at a 40% recall cutoff (N=224), the precision versus recall of the top 10 ECs is plotted. ECs are considered true if inter-residue minimum atom > 8Å, and false if > 12Å. (C)

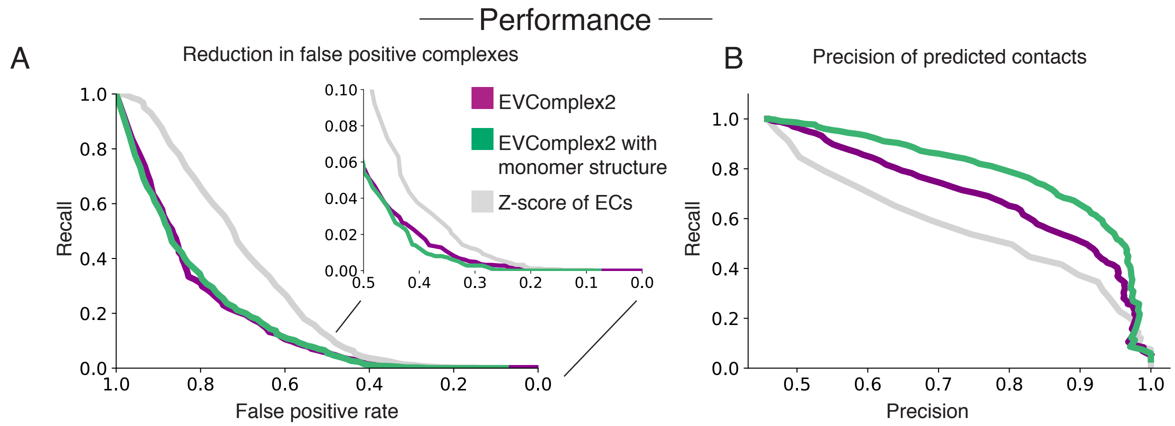

**Supplemental Figure 3: Protein interaction scores for cell envelope proteome.** Orange dots indicate proteins with a solved crystal structure of their interaction, grey dots indicate all other proteins. Protein pairs are plotted with the protein interaction score (X-axis) and the Z-score of their top inter-protein EC.

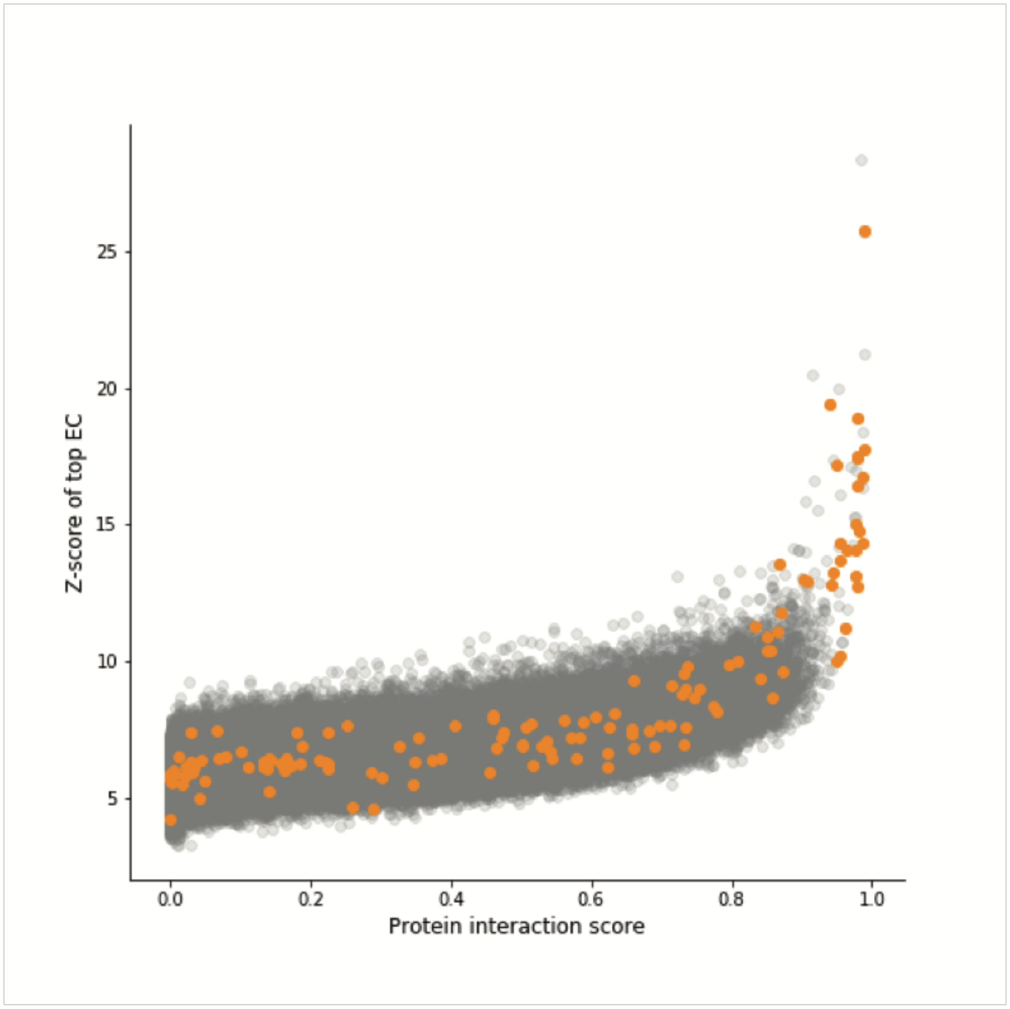

**Supplementary Figure 4: Inferred ring structure of flagellar hook-filament junction from three species**

Ring structure of FlgK (blue) and FlgL (green) from three different species (A) *Campylobacter jejuni,* (B) *Salmonella typhimurium,* (C) *Escherichia coli*. The *S. typhimurium* ring model was constructed by docking monomer structures 2D4Y and 2D4X using evolutionary couplings, and then arranging eleven copies of the lowest energy model in a ring based on the coordinates of the *C. jejuni* FlgK ring model[1]. The *E. coli* ring model was created by making a homology model[2] of the *E. coli* FlgL and FlgK proteins against the monomer structures 2D4X and 2D4Y, performing docking using evolutionary couplings, and then arranging eleven copies of the lowest energy model in a ring based on the coordinates of the *C. jejuni* FlgK ring model. The *C. jejuni* ring was created using the structure 5XBJ of FlgK and a homology model of the *C. jejuni* FlgL created with 2D4X as a template, performing docking using evolutionary couplings, and then arranging eleven copies of the lowest energy model in a ring based on the coordinates of the *C. jejuni* FlgK ring model.

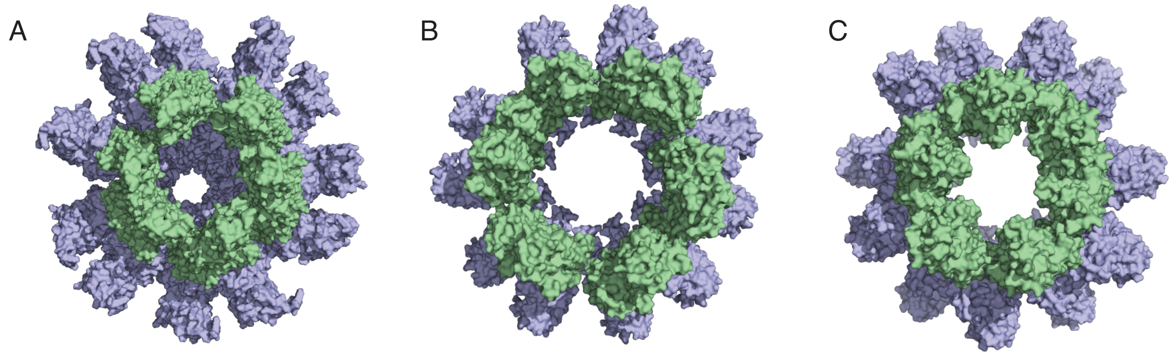
